# Negative trade-off between neoantigen repertoire breadth and the specificity of HLA-I molecules shapes antitumour immunity

**DOI:** 10.1101/2020.06.25.170472

**Authors:** Máté Manczinger, Gergő Balogh, Benjamin Tamás Papp, Balázs Koncz, Leó Asztalos, Lajos Kemény, Balázs Papp, Csaba Pál

**Author notes:** Correspondence should be addressed to MM or CP. These authors have contributed equally to this work (Máté Manczinger, Gergő Balogh, Benjamin Tamás Papp).

## Abstract

The human leukocyte antigen class I (*HLA*-I) genes shape our immune response against pathogens and cancer. Certain HLA-I variants can bind a much wider range of peptides than others, a feature that could be favorable against a range of viral diseases. However, the implications of this phenomenon on cancer immune response is unknown. In this paper, we quantified peptide repertoire breadth (or promiscuity) of a representative set of HLA-I alleles, and found that cancer patients that carry HLA-I alleles with high peptide binding promiscuity are characterized by significantly worse prognosis after immune checkpoint inhibitor treatment. This trend can be explained by a reduced capacity of promiscuous HLA-I molecules to discriminate between human self and tumour peptides, yielding a shift in regulation of T-cells in the tumour microenvironment from activation to tolerance. In summary, HLA-I peptide binding specificity shapes neopeptide immunogenicity and the self-immunopeptidome repertoire in an antagonistic manner. It could also underlie a negative trade-off between antitumour immunity and the genetic susceptibility to viral infections.

## Introduction

HLA-I molecules generally reside on the surface of human nucleated cells, and present intracellularly derived peptides. HLA-I genes are the most variable genes in the human genome^1^, and this variation shapes susceptibility to infectious diseases, autoimmune disorders and cancers^1^.

The adaptive immune recognition of mutated peptides (neopeptides) is essential for an effective tumor destruction by effector immune cells^2^. These neopeptides are presented on the surface of cancer cells generally bound by HLA-I molecules, and are subsequently recognized by T cells which can potentially induce an anticancer immune response^3^. Allelic HLA-I variation shapes the efficacy of antitumor immunity in a complex manner. HLA-I homozygosity, certain HLA-I supertypes and germline HLA-I evolutionary divergence (HED) are associated with an altered overall survival in patients receiving immunotherapy^4,5^. Additionally, HLA-I genotype shapes the mutational landscape of tumors: common missense mutations are generally bound weaker by HLA molecules^6^ and lung cancer cells frequently lose one (or both) of their HLA-I alleles, resulting in immune escape of cancer cells^7^.

An important feature of HLA-I alleles that has received little attention in cancer immunity is peptide binding promiscuity. There is a substantial variation in the size of the bound and presented peptide repertoire across HLA-I alleles^8–10^. Certain HLA-I alleles are capable of binding an exceptionally large set of peptide segments. For instance, a bioinformatics analysis has revealed over 16-fold variation in the number of peptides bound by common HLA-I alleles from a wide-range of dengue virus–derived peptides^10^. Similarly, the binding capacity of HLA-B molecules to a large set of self-peptides varies extensively, and influences the native repertoire of T-cell clones developing in the thymus^11^. Previous studies suggest that HLA class I and II peptide binding promiscuity has been positively selected during the evolutionary diversification of human populations, as they provide an enhanced capacity to withstand high pathogen diversity^8,9^.

In this work, we study the impact of peptide-binding repertoire of HLA-I alleles on cancer immune checkpoint inhibitor (ICI) therapy. ICIs target inhibitory receptors on T cells and thereby stimulate antitumour immunity^12^. However, only a subset of patients responds adequately to ICI based immunotherapies. Therefore, understanding the immunological mechanisms underlying therapeutic success is of paramount importance. We found that cancer patients carrying HLA-I alleles with promiscuous peptide binding are characterized by significantly *worse* prognosis after ICI immunotherapy. This trend can be explained by a reduced capacity of highly promiscuous HLA-I variants to discriminate tumour neopeptides from self-peptides. As a consequence, cancer samples from patients with promiscuous HLA-I alleles display signatures of an immunosuppressive tumour microenvironment with prominent T-cell dysfunction. Taken together, these results indicate that HLA-I alleles with low peptide binding specificities represent a genetic barrier to effective cancer immunotherapy.

## Results

### Peptide binding specificity of HLA-I alleles

Our first aim was to estimate the peptide-binding specificity of individual HLA-I alleles by collecting peptide–HLA-I interactions validated experimentally in vitro (see Methods). In total, we analyzed over 250000 peptide-HLA interactions covering 67 HLA-A, -B or -C alleles with appropriate *in vitro* data. These alleles are present at detectable frequencies and cover a large fraction of individuals in numerous human populations. Importantly, all HLA-A and HLA-B alleles of a well-established reference set with maximal population coverage^13^ were included in our analysis.

An established protocol was used to identify the amino acid composition of the peptides bound by the peptide binding domain of each HLA-I allele (see Methods). Using these data, we calculated peptide binding promiscuity (Pr), an index that estimates the sequence diversity of the peptides presented by each allele (see Methods). Alleles with high Pr are capable of binding a wide-variety of peptides and can be explained with diverse peptide motifs (Supplementary Fig 1). Bound peptides are enriched in either hydrophobic, basic amino acids or in tyrosine at anchor positions 2 and/or 9 (Supplementary Fig 1). At the same time, the selectivity at these and other positions are low compared to other alleles (Supplementary Fig 1).

We found a substantial variation in Pr across common HLA-A, HLA-B and HLA-C alleles (Figure 1A). The reliability of the calculated Pr values was first confirmed by analyzing the data of two large-scale immunopeptidomics studies^14,15^. These studies identified naturally eluted self-peptides from the surface of HLA-I monoallelic cell lines, and include peptide data for 10 HLA-A, 6 HLA-B and 15 HLA-C alleles. We found a strong positive correlation between Pr and peptidome diversity on the cell surface (Figure 1B). The correlation remained when data of the two studies were analyzed separately (Supplementary Fig 2).

**Figure 1.**
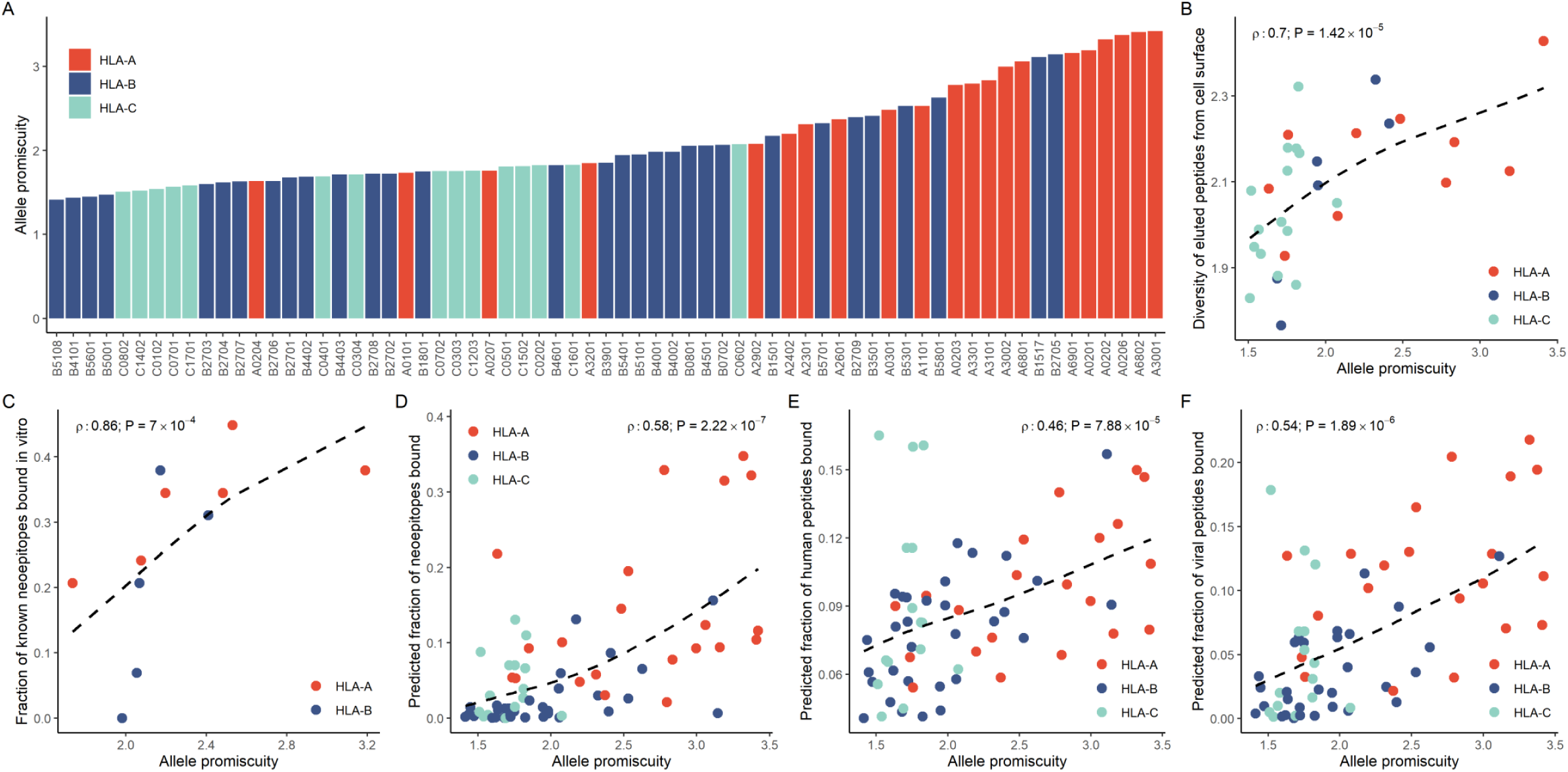
Basic properties of HLA-I allelic promiscuity. A) The promiscuity (Pr) values for the 67 HLA class I alleles used in our study are shown in increasing order. B) Amino acid diversity of self-peptides presented by the given allele on monoallelic cell lines is shown as the function of allele promiscuity. Alleles with higher promiscuity present more diverse peptide sequences C) The fraction of immunogenic neoepitopes bound by the given allele is shown as the function of allele promiscuity. Binding was determined in vitro with ProImmune REVEAL assays. D-F) The predicted fraction of bound neopeptides (D), human-peptides identified in immunopeptidomic studies (E) and viral peptides (F) is shown as the function of HLA promiscuity. On plots B to E, Spearman’s coefficient is shown with the level of significance and dashed lines indicate smooth curve fitted using cubic smoothing spline method in R (see Methods).

Additionally, the *in vitro* binding affinities of 11 representative HLA-I alleles to 29 tumour neoepitopes were measured with ProImmune REVEAL HLA-peptide binding assays. These alleles are widespread, and are consequently part of a small set of reference HLA variants that allows maximal covering of human populations^13^. The selected peptides were collected from the TANTIGEN database^16^. These peptides elicit a cytotoxic immune response (Supplementary Data), and display limited or no overlap in sequence (see Methods). The analysis revealed a substantial variation in the binding affinities to different peptides (Supplementary Data). Reassuringly, there was a strong positive correlation between Pr and the fraction of neopeptides bound by a given allele (Figure 1C, Spearman’s rho: 0.86, P = 7*10^-4^).

To confirm the results reported above, the binding specificities of HLA-I alleles against established sets of 1929 cancer peptides (Supplementary Data) and 9544 viral peptides derived from diverse pathogens (Supplementary Data) were calculated using the NetMHCpan-4.0 algorithm^17^. Additionally, it was also tested whether Pr is associated with the diversity of human self-immunopeptidome, using a set of 212,090 peptides identified in 25 immunopeptidomics studies (Supplementary Data). It was found that HLA-I variants with high Pr present a larger diversity of neopeptides, human self-peptides and viral peptides (Figure 1, D to F, Supplementary Data). These associations remained significant in multivariate linear regression models including the HLA-I locus as a categorical predictor variable (Supplementary Table 1). Together, these results indicate that HLA-I peptide binding promiscuity shapes the diversity of presented tumour, human self and viral immunopeptidome as well. They also verify that Pr is a suitable index for estimating the diversity of peptides presented by HLA-I alleles.

As the stability of peptide-HLA complexes is essential for an effective antigen presentation^18^, we finally evaluated whether promiscuous HLA-I alleles have an altered capacity to form stable protein complexes with neoepitopes. The assembly and dissociation rates of 66 representative neoepitope-HLA-I complexes were measured in vitro using ProImmune Complete rate assays (see Methods). On average, the assembly of the complexes tends to be faster and the resulting complexes are more stable with high Pr HLA-I molecules (Figure 2A to D). These results were further supported by the computational analysis of HLA-peptide complex stability, involving 67 HLA alleles and 1929 neopeptides (Figure 2E, see Methods for details). We conclude that, in contrast to other proteins^19^, there is no evidence for a negative trade-off between binding multispecificity and complex stability in case of HLA-I molecules.

**Figure 2.**
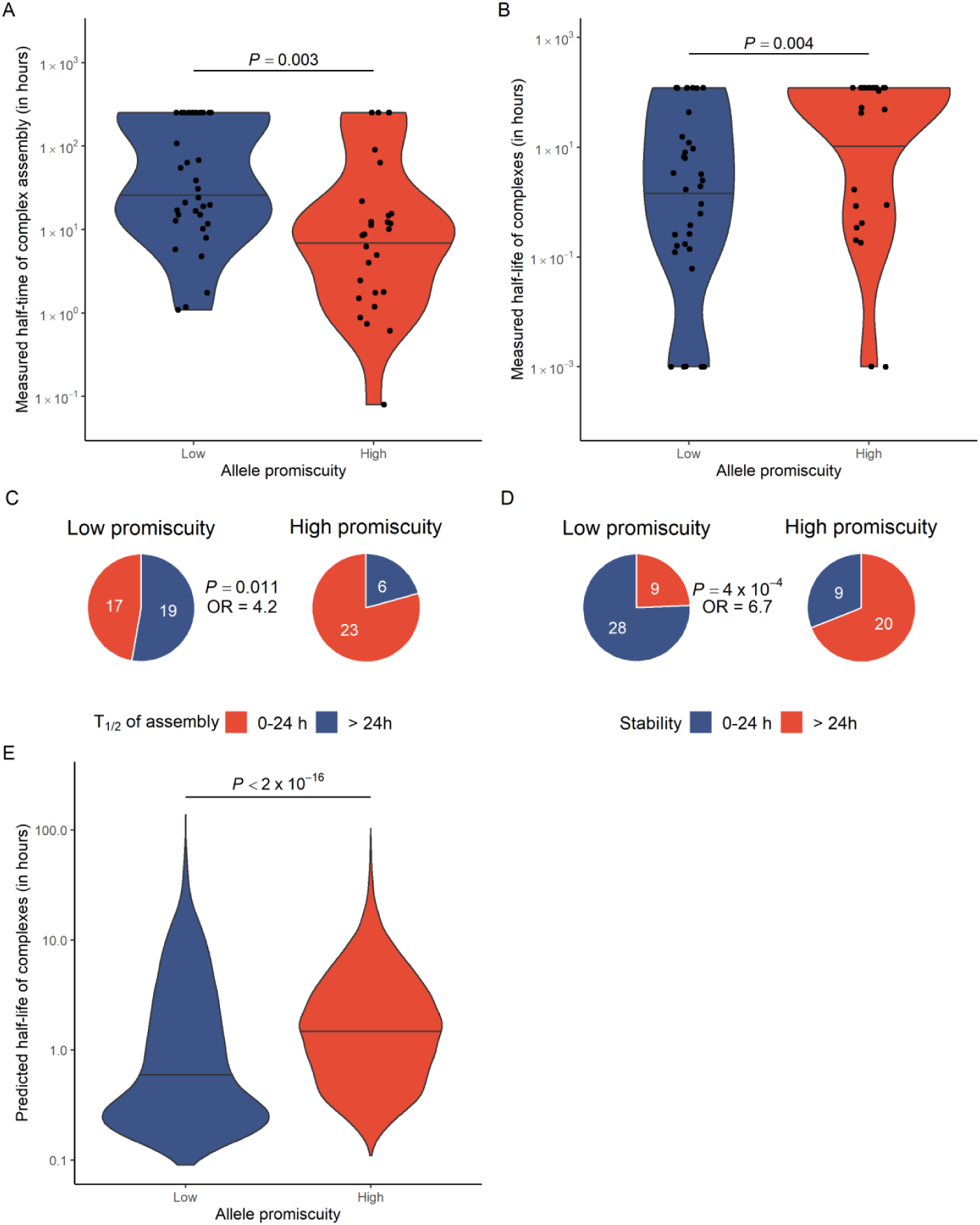
High allele promiscuity is associated with more stable peptide-HLA complexes. A) In vitro measured half-time of complex assembly for 66 allele-neoepitope pairs. Alleles were stratified into low and high promiscuity groups based on median. B) Peptide-HLA complexes of promiscuous HLA variants are more stable. The Y axis shows complex half-life in hours. C) Promiscuous alleles are 4.2-times more likely to form complexes with an assembly half-time less than 24 hours and D) 6.7-times more likely to form especially stable complexes with a half-life longer than 24 hours. For panel C and D, the P-values for Fisher’s exact tests are indicated. E) Predicted half-life of neopeptide-HLA complexes, using the NetMHCstabpan algorithm^23^. Complexes were identified by predicting the binding affinity of 1929 neopeptides to 67 HLA-I alleles (see Figure 1D and Methods). Alleles were stratified into low and high promiscuity groups using the same cutoff as for panel A and B. On panels A, B and E, the P-value for Wilcoxon’s rank-sum tests are indicated. Violin plots show the density function of values indicated on Y axes and horizontal lines indicate the median value in each group.

### High level of HLA-I promiscuity yields worse prognosis

To investigate the impact of Pr on cancer immunotherapy, we focused on previously published cohorts of cancer patients treated with ICI^4,20–22^. We collected data on melanoma patients receiving anti-CTLA-4 therapy^4,20,21^ (N = 164), NSCLC patients treated with anti-PD-1 therapy^22^ (N = 74), and melanoma patients treated with anti-PD-1/anti-PDL-1 therapy^4^ (N = 89). The HLA genotype of patients in these three cohorts were determined previously^4^. For each patient, we calculated genotype Pr, as the mean of six HLA-A, HLA-B and HLA-C allelic Pr values, under the assumption that that each locus contributes to the presentation of neopeptides. There was a substantial variation in genotype Pr across patients in the examined cohorts (Supplementary Fig. 3).

We next examined how genotype Pr affects overall survival probability in response to ICI therapy. Patients were classified into high, medium and low genotype Pr, based on the top and bottom quartiles as cutoffs. High genotype Pr was found to be associated with reduced overall survival in all three cohorts. (Figure 3A to C). For example, survival rates of high genotype Pr melanoma patients treated with CTLA-4 inhibitors were 54% lower than that of patients in the low Pr group, while the high genotype Pr group of NSCLC patients treated with PD-1 inhibitors were 57% less likely to survive than individuals in the low Pr group.

**Figure 3.**
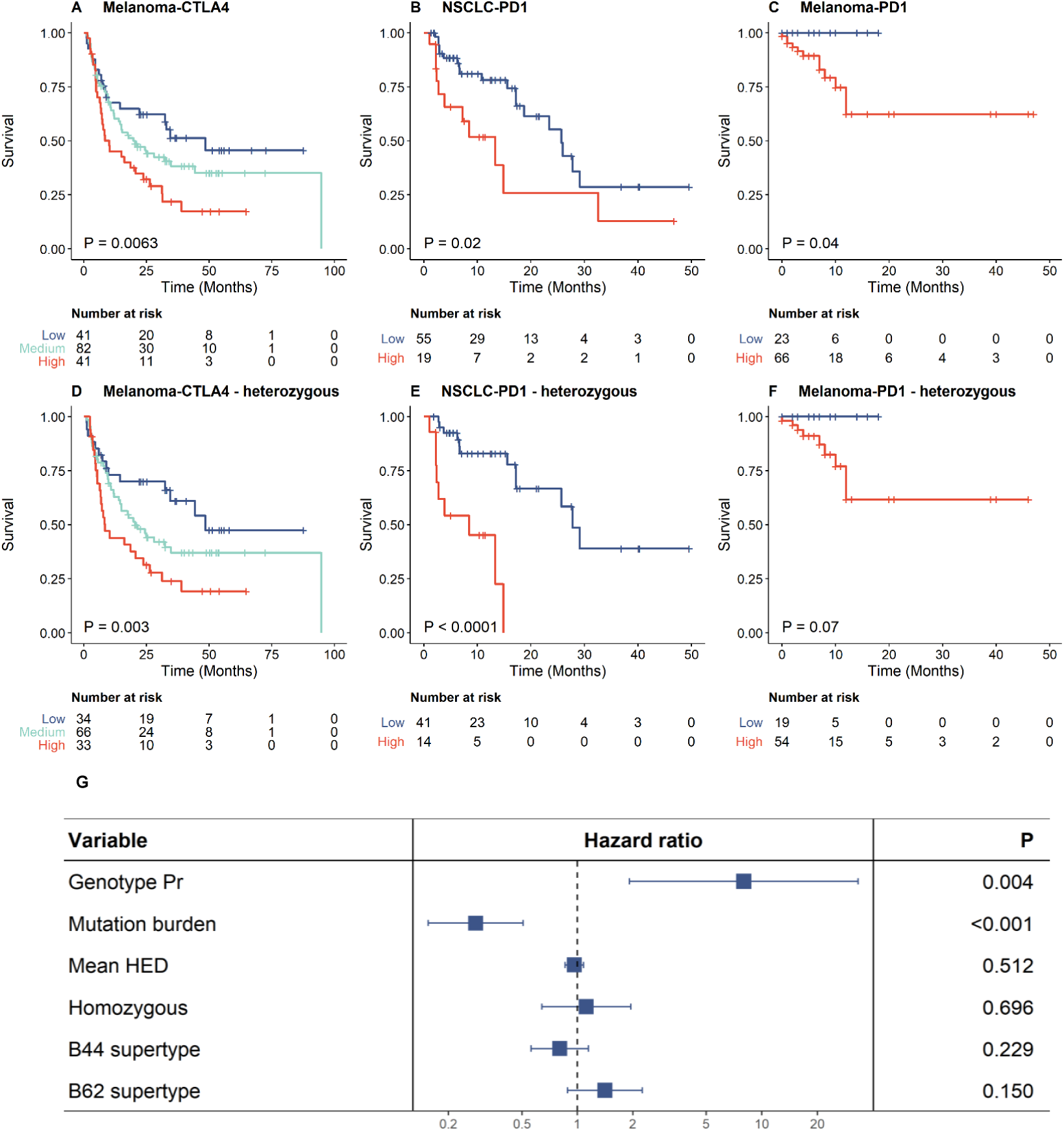
High allele promiscuity and survival rate of melanoma patients receiving anti-CTLA-4 therapy (A, D), NSCLC patients receiving anti-PD1 therapy (B, E) and melanoma patients receiving anti-PD1/anti-PD-L1 therapy (C, F). Patients were classified into two (B-C and E-F) or three (A and D) groups based on quartiles of genotype Pr. Promiscuity was cut at the upper quartile for B and E and at the lower quartile for C and F. Log-rank test P-values are shown. Between-group differences were also tested for trend in case of A and B (see Methods). The analysis was carried out for all patients (A-C) or only for patients fully heterozygous at the HLA-I loci (D-F). G) The effect of genotype Pr on survival remained highly significant in a multiple Cox regression model including mutation burden and other HLA-associated features as covariates. The model involved all patients from panel A to C. Genotype Pr, mutation burden and mean HED were treated as continuous variables.

### HLA-I promiscuity is a major determinant of patient survival

The relationship between the level of HLA-I promiscuity and survival remained significant or became even stronger when patients homozygous on any of the HLA-I loci were excluded from the analysis (Figure 3, D to F). Similarly, the association remained significant when the analysis was iteratively repeated by excluding individuals carrying specific HLA alleles (Supplementary Table 2). Hence, the impact of HLA promiscuity on patient survival cannot be explained by a single, highly prevalent allele with peculiar peptide binding properties.

Based on the structural properties of the peptide binding regions, genetically related HLA-I alleles were previously classified into 12 supertypes that cover the majority of common HLA-A and HLA-B alleles^24^. Melanoma patients carrying HLA-B alleles belonging to the B44 superfamily were reported to have significantly better survival after ICI treatment, but the reasons have remained unclear^4^. Notably, the association between patient survival and B44 supertype is mainly due to specific HLA-B alleles, such as B*18:01, B*44:02, B*44:03, B*44:05, B*50:01. The four B44 alleles included in our analysis have particularly low Pr compared to other alleles (Supplementary Fig 4).

Tumor mutation burden^25^ (TMB), HLA-I heterozygosity^4^ and HLA-I evolutionary divergence^5^ (HED) have previously been suggested as the determinants of response to ICI therapy. These features are expected to increase the diversity of neopeptides that can be presented on the surface of tumour cells^12^. Genotype Pr showed no statistically significant relationship with HED and HLA-I heterozygosity (Supplementary Fig 5). Moreover, the negative association between genotype Pr and patient survival remained highly significant in a multiple Cox-regression model after controlling for these features (Figure 3G). In particular, TMB and genotype Pr appeared to be the strongest determinants, while HED and HLA-I heterozygosity had no significant effect on survival in this model. This is in line with a prior work showing no association between HLA class I heterozygosity and survival of NSCLC patients receiving immunotherapy^26^. Furthermore, the impact of Pr on patient survival remained after controlling for the effect of specific HLA-B supertypes B44 and B62, which have previously been associated with survival outcome^4^ (Figure 3G).

We next evaluated whether the low Pr of each HLA locus contributes to improved survival. For this purpose, the mean Pr was calculated for HLA-A, HLA-B and HLA-C loci individually and also for the combinations thereof. The promiscuity level of HLA-B was predictive of survival outcome, but the negative association between genotype Pr and patient survival becomes even stronger by inclusion of HLA-A and HLA-C into univariate Cox models (Supplementary Table 3). The prominent effect of HLA-B on patient survival might be explained by the relatively high abundance of this molecule on the cell surface relative to that of HLA-A and HLA-C^4,27^. Finally, genomic analysis of cancer mutations in 165 patients revealed no significant association between genotype Pr and the frequency of loss of heterozygosity events at any of the HLA-I loci (Wilcoxon’s rank-sum test P = 0.64).

### Reduced discrimination capacity of self and mutated peptides by promiscuous HLA-I molecules

We next asked whether the presentation of more diverse peptides by promiscuous HLA-I molecules results in a reduced capacity to differentiate between human self-peptides and their mutated counterparts (neopeptides). For this purpose, we calculated the differential agretopicity index^28^ (DAI), which is a broad indicator of neopeptide dissimilarity from self and thereby a feature of immunogenicity^28^. DAI estimates the difference between the predicted affinity for the mutant peptide and the corresponding non-mutated homologous peptide pair^28^. To gain insight into the capacity of each HLA-I molecule to discriminate between self and tumour peptides, we employed a set of 589 experimentally verified tumour neopeptides (see Methods). For each HLA-I variant, we calculated the median DAI for neopeptides bound by the given HLA-I allele (see Methods). A strong negative association was found between median DAI and the level of promiscuity for HLA-A and HLA-B alleles, but not for HLA-C alleles (Figure 4, A to C).

**Figure 4.**
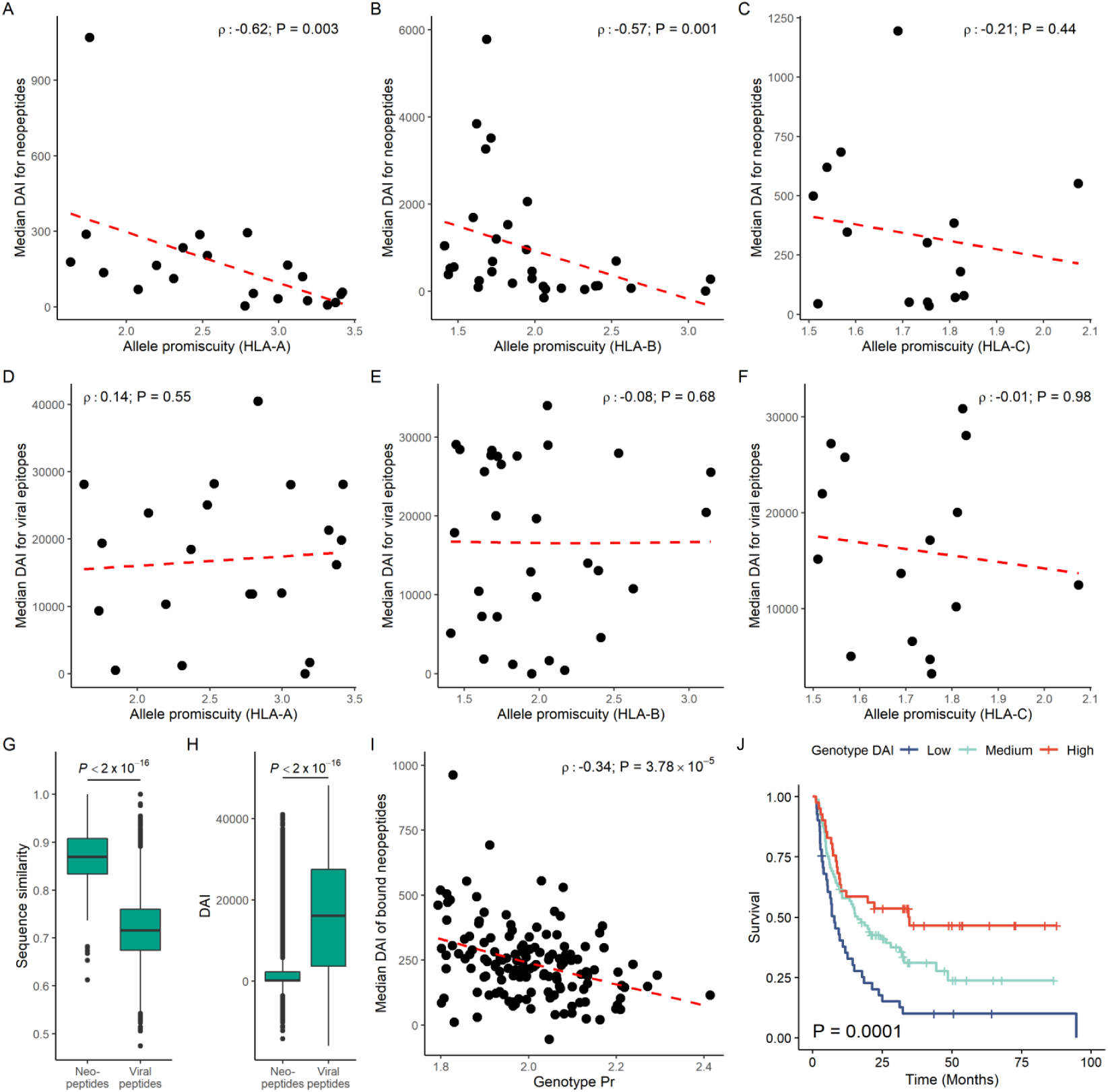
HLA-I allelic promiscuity and differentiation capacity between neopeptides and the corresponding non-mutated peptides. Relationship between the promiscuity of A) HLA-A, B) HLA-B and C) HLA-C alleles and median DAI for bound tumour neopeptides. Higher HLA-A and HLA-B promiscuity is associated with a lower median DAI value, while there is no significant relationship between the two variables in case of HLA-C. There is no statistically significant relationship between promiscuity of D) HLA-A, E) HLA-B and F) HLA-C alleles and the median DAI calculated for bound viral peptides. G) The maximum BLOSUM62 sequence similarity of neopeptides and viral peptides to the non-mutated human proteome. H) DAI values for HLA-bound viral peptides are much higher than for neopeptides. I) Bound neopeptides in melanoma samples with promiscuous HLA alleles have a lower median DAI. J) The same patients were stratified into low, medium and high genotype DAI groups based on the 1^st^ and 3^rd^ quartiles of all values. A higher median DAI was associated with better survival. Log-rank test P-value is shown. Between-group differences were also tested for trend (see Methods). Spearman’s rho and the associated P-values are shown on figures A to F and I. Wilcoxon’s rank sum test P-values are shown on figures G and H. On figures A to F and I, dashed lines indicate a smooth curve fitted using cubic smoothing spline method in R (see Methods). On boxplots, horizontal lines indicate median, boxes indicate the interquartile range (IQR), vertical lines indicate 1^st^ quartile - 1.5*IQR and 3^rd^ quartile + 1.5*IQR.

An elevated level of HLA-I promiscuity has been suggested to be favorable in terms of eliciting an immune response against diverse viral infections^8,9^. Therefore, it is important to establish whether the results above are specific to tumour neopeptides or they also hold for virus-derived epitopes. We analyzed a previously compiled set of 1030 viral epitopes with established positive T-cell assays^29^. To calculate DAI for viral peptides, each peptide was aligned against the non-mutated human proteome and the closest human hit was retrieved to calculate relative peptide-HLA-I binding affinities, as described earlier^30^. The analysis revealed no significant association between median DAI of bound viral peptides and allelic HLA-I promiscuity level (Figure 4, D to F). As most studied viral peptides displayed only a limited sequence similarity to the human proteome (Figure 4G), DAI values for viral peptides were much higher than for neopeptides (Figure 4H). Consequently, even HLA molecules with a low peptide binding specificity could be able to discriminate between viral and human-self peptides.

It has been previously suggested that tumours carrying high DAI neopeptides are more susceptible to immune recognition and hence should be responsive to ICI therapy^28,31–33^. Therefore, a next important question is how HLA-I promiscuity shapes the abundance of potentially immunogenic, high DAI neopeptides in cancer patients. For this purpose, we analyzed the cancer genomes of 139 melanoma patients treated with CTLA-4 inhibitors^20,21^. From the detected set of non-synonymous mutations in each cancer sample, we determined the complete set of neopeptides bound to at least one of the HLA-I alleles in the corresponding patient (see Methods). The median DAI for neopeptides showed a significant negative association with genotype Pr (Figure 4I) indicating that that the immunogenicity of neopeptides is contingent upon HLA-I promiscuity level. As earlier^28,30^, we also found that higher median DAI is associated positively with patient survival (Figure 4J). These results do not simply reflect differences in the set of cancer mutations across patients. For each genotype, DAI was calculated as the mean of allele-specific values determined on a fixed set of 589 experimentally confirmed neopeptides, and the positive association between DAI and survival was found to remain (Supplementary Fig 6). Together, these results indicate that patients who carry promiscuous HLA-I alleles have a reduced capacity to differentiate between self-peptides and the corresponding mutant tumour neopeptides with implications on patient survival^28^.

### HLA-I promiscuity and T-cell tolerance

Immunogenic neopeptides need to have a strong binding affinity to HLA-I/T-cell receptor complexes. However, when their non-mutated counterparts are also presented by HLA molecules, the corresponding CD8^+^ T-cells reactive to the neopeptides are expected to be removed by central tolerance in the thymus or are inhibited by peripheral tolerance^28,33^. Here we focus on the impact of HLA-I promiscuity on peripheral tolerance, a mechanism that ensures that self-reactive T and B cells which have not been eliminated by central tolerance in the thymus do not recognize self-peptides as foreign ones and initiate auto-immune reactions^34^. However, elevated level of T-cell tolerance in the tumour microenvironment contributes to uncontrolled tumor growth^35,36^. We hypothesized that due to the reduced discrimination capacity of self and tumour peptides in patients carrying high Pr HLA-I alleles, regulation of T-cells in the tumour microenvironment is shifted from activation to tolerance, yielding reduced capacity to eliminate cancer cells upon ICI therapy.

To identify molecular and cellular signatures of T-cell tolerance in cancer samples, we focused on datasets from melanoma patients treated with the anti-CTLA4 antibody ipilimumab^20^. A unique aspect of this dataset is that it combines information on the transcriptome for a sufficient number of pre-treatment cancer samples with HLA-genotype, and disease progression of each patient.

We first performed a gene set enrichment analysis comparing patients with high and low genotype Pr. Most notably, the genes involved in the positive regulation of T-cell tolerance induction, type 2 immune response, extracellular matrix secretion and macrophage induction were found to be up-regulated in high Pr cancer samples (Figure 5A). Similarly, the genes associated with the negative regulation of T helper 1 cell mediated immune response and CD4^+^ alpha beta T-cell differentiation were expressed at higher levels in high Pr samples (Figure 5A). These cellular processes are linked to the induction of an immunosuppressive tumor microenviroment^36,37^. Specifically, macrophages inhibit effector immune responses and stimulate angiogenesis^38^. Additionally, they skew type 1 immune response (mediated by T helper 1 cells) towards type 2, which is less effective against cancer cells^37,38^. Remodeling of the extracellular matrix is essential for immune suppression and cancer progression^39,40^. Finally, the lower level of CD4^+^ T-cell differentiation and T-cell tolerance induction prohibit effective tumor destruction^35,36^.

**Figure 5.**
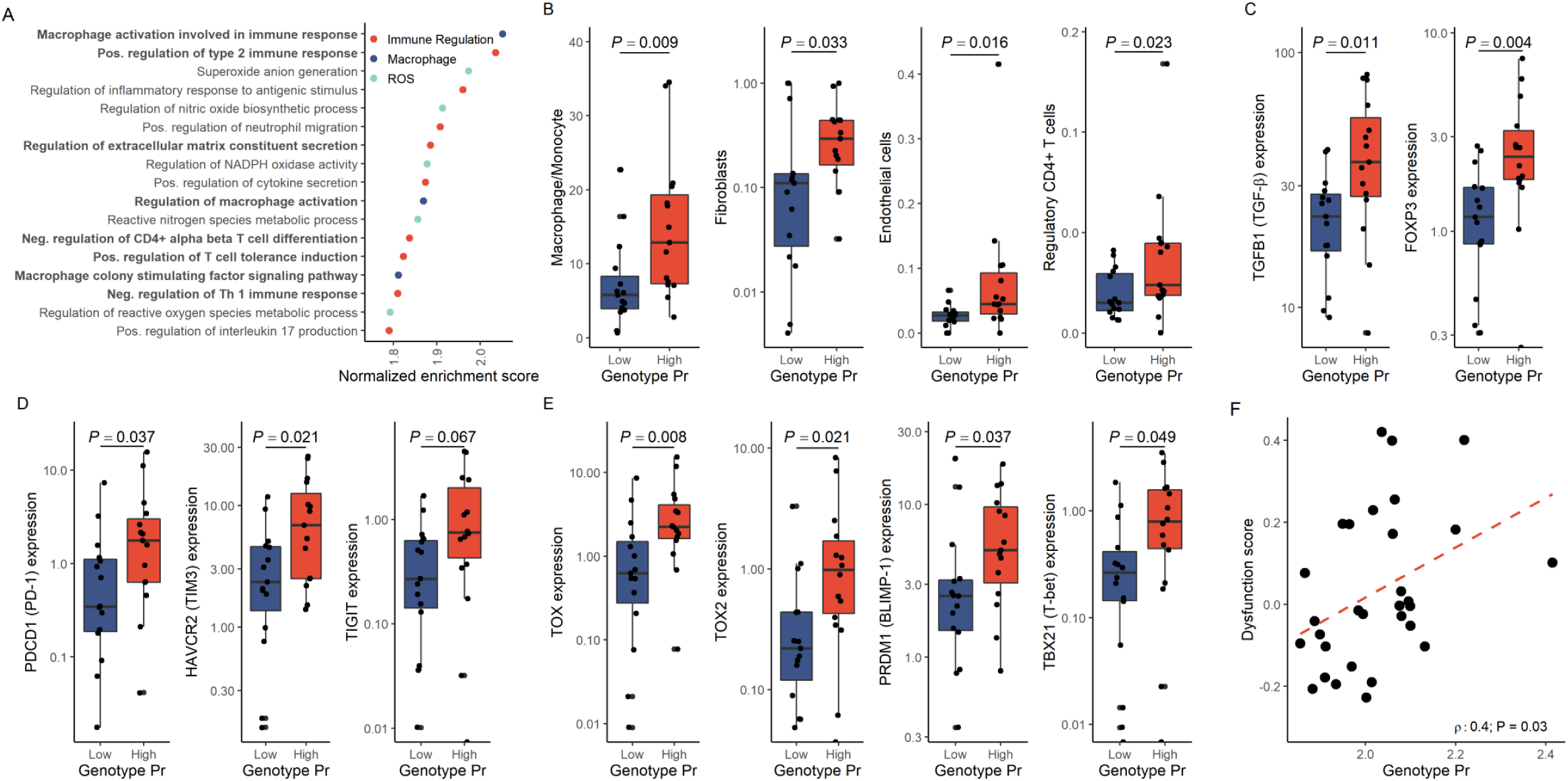
Association of high HLA genotype promiscuity with an immunosuppressive cancer microenvironment. Patiens were stratified into low and high genotype Pr groups using the median as a cutoff. A) Results of gene set enrichment analysis. Gene sets enriched in high genotype Pr samples with a P < 0.001 and FDR < 0.1 are shown. Terms are classified into three main categories. Terms associated with the development of an immunosuppressive tumor microenvironment are highlighted in bold. B) Results of immune deconvolution analysis. Boxplots show approximated cell abundance in samples with low and high genotype Pr is shown on boxplots. C) Expression of genes encoding the cytokine TGF-β and the transcription factor FOXP3, molecules associated with T-cell tolerance. D) Expression levels of genes encoding checkpoint molecules are indicated on the figure. E) Expression of genes encoding master regulators associated with T cell dysfunction. F) High genotype Pr is associated with and elevated T-cell dysfunction score calculated by the TIDE algorithm. Spearman’s rho and correlation P-values are indicated. Dashed red line indicates linear regression line. On panels B to E, P-values for Wilcoxon’s rank sum tests are shown. On boxplots, horizontal lines indicate median, boxes indicate the interquartile range, vertical lines indicate 1^st^ quartile - 1.5*IQR and 3^rd^ quartile + 1.5*IQR.

Next, by applying state-of-the-art immune deconvolution methods and recommendations of prior benchmark studies^41^, we assessed the immune cell composition in these tumor samples^41^. The analysis revealed an overrepresentation of cells associated with immunosuppression in the tumour samples derived from patients with high genotype Pr (Figure 5B), including regulatory T-cells (Tregs), cancer associated fibroblasts, macrophages/monocytes and endothelial cells. Tregs and macrophages produce immunosuppressive mediators, such as TGF-β, which downregulates effector immune functions, and induce the development and survival of regulatory T-cells^42^. As expected, cancer samples from high Pr patients displayed increased expression of TGFB1 encoding TGF-β (Figure 5C). This is significant, as elevated TGF-β expression is associated with the promotion of tumorigenesis^43^. Similarly, FOXP3 gene - that encodes a master regulator protein involved in the development and function of Tregs – was also highly expressed in these samples^35^ (Figure 5C).

T-cell co-stimulation is an essential step in developing T-cell tolerance or function^35^. Negative costimulatory molecules inhibit T-cell activation at immune checkpoints leading to T-cell tolerance^35^. Consequently, the inhibitory immune checkpoint receptor PD-1, as well as the and co-inhibitory receptors TIGIT and TIM3 have established roles in promoting tumor immune evasion^44,45^. Genes encoding these three molecules displayed significantly elevated expression in cancer samples with high Pr (Figure 5D). It is well-known that TIM3 is especially abundant in FOXP3 expressing Treg cells and macrophages^46^.

Irreversible dysfunction of T-cells is a common cause of non-responsiveness to immune checkpoint blockade therapy^47^, and is linked to the formation of an immunosuppressive tumor microenvironment^36^. Therefore, we next asked whether high Pr HLA-I alleles are associated with signatures of T-cell dysfunction. Remarkably, we found that cancer samples with high Pr display high expresson levels of genes encoding TOX, TOX2, T-bet and BLIMP1 (Figure 5E). These transcription factors are critical for the progression and the maintenance of T-cell dysfunction^48,49^. To investigate T-cell dysfunction signatures in cancer samples more systematically, we employed the Tumor Immune Dysfunction and Exclusion (TIDE) method^47^. TIDE is a recently developed computational tool providing genome-wide scores of T-cell dysfunction and T-cell exclusion signatures for each tumor and can reliably predict ICI response. Importantly, the dysfunction score describes the level of late-stage, irreversible dysfunction of T-cells as reported by the authors^47^. We found a positive correlation between genotype Pr and T-cell dysfunction score (Figure 5F, Spearman’s rho: 0.4, P = 0.03), but not between genotype Pr and T-cell exclusion score (Spearman’s rho= -0.1, P = 0.6). This suggests that HLA-I promiscuity level does not affect T-cell infiltration directly, but rather it shapes the functioning of T-cells in the tumour microenvironment. Importantly, other clinically relevant HLA-I associated features, such as HLA-I heterozygosity, germ-line HLA-I evolutionary divergence and specific HLA-I supertypes showed no association with T-cell dysfunction score (Supplementary Fig 7).

Taken together, these results indicate that cancer samples carrying highly promiscuous HLA-I alleles tend to display signatures of immune-suppressive tumour microenvironment and T-cell dysfunction resulting in resistance to ICI therapy.

## Discussion

HLA-I genes encode a critical component of the adaptive immune system. The human genome contains six primary allelic variants encoded by HLA-A, HLB-B and HLA-C. Together, these alleles define the set of individual peptides presented to T-cells^1^. The resulting peptide repertoire – or immunopeptidome – substantially differs across individuals, due to the remarkable sequence variation in the peptide binding region of these genes. As a consequence, each allele presents a distinct, and only partially overlapping set of peptides. Moreover, HLA variants differ by orders of magnitude in the breadth of peptides they can bind^8,9^. HLA-I alleles with promiscuous peptide binding may be favorable in pathogen-rich environments^8^, but their role in anticancer immunity has remained unestablished.

To investigate this issue, we first developed a computational tool to systematically explore the sequence diversity of the peptide set presented by different HLA-I molecules, and found that high HLA-I promiscuity is associated with enhanced diversity of tumour, viral and self-immunopeptidomes as well (Figure 1 B, D to F). Next, we showed that melanoma and NSCLC cancer patients carrying HLA-I alleles with high Pr display significantly worse prognosis after ICI immunotherapy (Figure 3). This pattern is surprising, as the breadth of bound peptides by HLA-I molecules is expected to underlie a successful immunologic control of cancer. Therefore, we next aimed to establish the deleterious side-effects of elevated HLA-I promiscuity level on anti-cancer immunity.

The stability of the peptide–HLA-I complex is a strong predictor of antigen immunogenicity^18^. In contrast to expectations, promiscuous HLA variants tend to form more stable HLA-peptide complexes (Figure 2). This finding has important immunological implications and should be further investigated in future research. On the other hand, we found a strong negative association between the level of promiscuity and the differential binding affinity of mutated neopeptides (Figure 4). This suggests that HLA variants with promiscuous peptide binding have a reduced capacity to discriminate self-peptides from mutated tumour peptides.

We hypothesized that as a consequence, regulation of T-cells in the tumour microenvironment is shifted from activation to tolerance, yielding reduced a capacity to eliminate cancer cells upon ICI therapy. In agreement with expectation, cancer samples from melanoma patients carrying promiscuous HLA-I alleles displayed molecular and cellular signatures of peripheral T-cell tolerance and an immunosuppressive tumour microenvironment (Figure 5). These signatures include increased prevalence of regulatory T-cells and other immunosuppressive cells in the tumour microenvironment, elevated expression of immune checkpoint inhibitory receptors (PD-1), suppressive soluble mediators (TGF-β) and master control genes involved in regulatory T-cell development and maintenance (e.g. FOXP3). Future work should elucidate the exact molecular pathways underlying these patterns. This research will require a series of HLA transgenic mice studies on the role of promiscuity in T-cell initiated immune response. The impact of HLA-I promiscuity level on central tolerance and potential depletion of T-cells in the repertoire in healthy individuals also remains to be elucidated^50^.

In sum, high peptide binding promiscuity could be a common feature of genetically diverse HLA-I variants associated with reduced survival upon ICI therapy. Therefore, this metric could potentially be used for future clinical trials to identify genetic variables that affect anti-tumor immunity. Importantly, HLA promiscuity is conceptually different from and statistically independent of other, previously identified determinants driving response and adverse effects of ICI immunotherapy^4,5,12^ (Supplementary Fig 5).

Our work has several important implications for future studies. First, HLA class II genotype is known to drive anti-tumor responses and tumour evolution in general^51,52^. Therefore, it is important to establish how the functional diversity of HLA-II molecules shapes the efficacy of ICI immunotherapy. Second, while the presentation of a broad range of viral peptides by promiscuous HLA-I alleles could be favorable in terms of acute viral infections^8,9^, our work indicates that such alleles impede anti-tumour immunity. In other words, Pr could underlie a negative trade-off between antitumour immunity and the genetic susceptibility to viral infections. Why is it so? As viral peptides tend to display limited sequence similarity to the human proteome^30,53^ (Figure 4G), the limited discrimination capacity between self and non-self-peptides by promiscuous HLA alleles may be less of an issue in the case of antiviral immune responses (Figure 4H). Consistent with this hypothesis, we found no correlation between the level of HLA-I promiscuity and differential binding affinity of viral epitopes and self-peptides across the non-mutated human proteome (Figure 4, D to F). Therefore, promiscuous HLA-I molecules present a wider-range of viral epitopes (Figure 1F) without paying the cost of reduced discrimination between self and virus derived peptides. Third, and on a related note, promiscuous HLA-A and HLA-B variants are especially wide-spread in South-East Asia, possibly as a result of selection to withstand high pathogen load^8^. This pattern raises the possibility that there are geographical differences in the genetic make-up shaping the efficacy of cancer immunotherapy and anti-tumour immunity in general. Finally, prior studies suggested that neoantigen immunogenicity and quality, and not simply neoantigen quantity that determine patient survival^54^. Our work indicates that HLA-I peptide binding promiscuity shapes neoantigen immunogenicity with implications on the design of neoantigen vaccines^55^.

## Acknowledgments

We thank Tobias Lenz for earlier discussions on this topic, and providing us with the raw data on human evolutionary divergence (HED) values of HLA-I allele pairs. We are also grateful for Balázs Győrffy for his suggestions in survival analysis. The study was supported by the following research grants: The European Research Council H2020-ERC-2014-CoG 648364 – Resistance Evolution (C.P.) ‘Célzott Lendület’ Programme of the Hungarian Academy of Sciences LP-2017–10/2017 (CP); ‘Élvonal’ KKP 126506 (C.P.), GINOP-2.3.2–15–2016–00014 (EVOMER, for C.P. and B.P.), GINOP-2.3.2– 15–2016–00020 (MolMedEx TUMORDNS for C.P.), GINOP-2.3.3–15–2016–00001, GINOP-2.3.2–15– 2016–00026 (for B.P.), National Research, Development and Innovation Office (FK 128775 to ZF and KKP 129814 to B.P.) and The European Union’s Horizon 2020 research and innovation programme under grant agreement No 739593 (B.P). M.M. was supported by the ÚNKP-19-4 New National Excellence Program of the Ministry of Human Capacities, and by the Bolyai János Research Fellowship of the Hungarian Academy of Sciences. L.A. was supported by the ÚNKP-19-2 New National Excellence Program of the Ministry for Innovation and Technology. G.B. was supported by the ÚNKP-19-3 New National Excellence Program of the Ministry for Innovation and Technology. The authors thank Dora Bokor, PharmD, for proofreading the manuscript.

## Methods

### Calculating peptide binding specificity

Data on peptide-HLA class I interactions and corresponding T-cell activation assays were downloaded from The Immune Epitope Database^1^ (IEDB), as of 17^th^ January 2019. Only peptides with lengths between 8 to 12 standard amino acids, and with positive MHC binding and/or T-cell activation assays were considered further. To ensure reliable estimates on peptide binding promiscuity, only HLA-I alleles with data on the binding of at least 400 different peptides were analyzed further. This procedure resulted in a total of 253,147 peptide-HLA I interactions across 21 HLA-A, 31 HLA-B, and 15 HLA-C alleles. HLA-I allelic promiscuity was calculated as follows. Peptides were classified into discrete peptide sets according to their amino acid lengths and each set was analyzed separately. At each amino acid position, the amino acid frequency distribution in the peptide set was compared with the amino acid frequency distribution in the complete human proteome (downloaded from UniProt^2^), using Kullback-Leibler (KL) divergence. KL divergence is a standard metric regularly used to measure the distance between two discrete probability distributions (*P* and *Q*) and is calculated as follows:

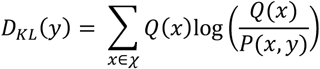

where *P(x,y)* denotes the frequency of amino acid *x* in the peptide set at amino acid position *y*, while *Q(x)* denotes the frequency of the same amino acid in the complete human proteome (*χ* is the set of 20 amino acids). Position-specific D_KL_(y) values were calculated by the Rtreemix R library^3^. Finally, position specific D_KL_(y) values were averaged across the full peptide length and for each peptide set, yielding 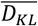. The more selective the binding of HLA-I allele is at a given position the higher the deviation is between the two distributions, resulting in higher 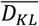. To control for any potential sample size associated biases, the raw 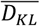 values were normalized by subtracting 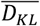 (rand). 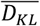 (rand) is the average of 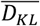 values based on the same number of randomly selected peptides with the same length in the human proteome iterated 1000 times. The normalized 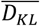 values show no significant correlation with the number of input peptide sequences (Spearman’s rho: -0.08, P = 0.23). HLA allelic promiscuity (Pr) is defined as the reciprocal value of normalized 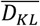 i.e. 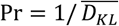 Pr values do not change significantly when using *Q(x)* derived from the proteomes of bacterial or viral species instead of the complete human proteome (Supplementary Fig 8C and D). Computer simulations also showed that Pr can be reliably inferred using the above defined threshold of peptide number per HLA allele (Supplementary Fig 8A and B). Of note, Kullback-Leibler divergence, which measures the discrepancy between two probability distributions is conceptually related to the Shannon entropy index^4^. As expected, the Pr values based on Shannon entropy index and Kullback-Leibler divergence show strong positive correlation with each other (Spearman’s rho: 0.86, P < 2.2*10-16).

### Experimental analysis of HLA-I neoepitope binding affinity

HLA-I peptide neoepitopes with established roles in mediating antitumor immune responses were collected from the TANTIGEN database^5^ (Supplementary Data). Epitopes with over 50% sequence identity were excluded using an iterative method^6^, resulting in 29 peptide neoepitopes. Each peptide was synthesized and their binding affinity towards a pre-selected set of 11 HLA-I variants was tested using the in vitro HLA class I Reveal binding assay (ProImmune Ltd., Oxford, UK). The binding assay determines the capability of each candidate peptide to bind to one or more HLA Class I alleles by assessing its ability to stabilize the HLA-peptide complex. For each allele, the binding of the tested peptides is scored relative to that of a corresponding high affinity T cell epitope.

### Experimental analysis of peptide-HLA complex assembly and stability

All peptide-HLA-I pairs with positive outcome of the HLA class I Reveal binding assay were studied further. The assembly and stability of 66 putative peptide-HLA complexes were measured using Complete Rate Assays (ProImmune Ltd., Oxford, UK). The fraction of assembled complexes was measured at six different time points over 48 hours. After fitting the values to a one-phase association equation, the half-time needed to reach the maximum of assembled complexes was calculated and reported as the on-rate of the studied peptide-HLA complex. The stability of the complexes was measured using off-rate assays. This assay measures the fraction of denaturated complexes at 6 different time points over 24 hours. These values were fitted to a one-phase denaturation equation and the half-life of complexes was calculated.

### Computational prediction of HLA binding affinity and complex stability

Using literature information^5,7–11^, we compiled datasets of i) 1929 experimentally verified neopeptides, all of which are known to have high binding affinity to certain HLA alleles (Supplementary data), ii) 212,090 HLA-I-bound self-peptides identified in a range of immunopeptidomics studies (Supplementary data) and iii) 9544 viral peptides acquired from a dataset that was published by Ogishi et al.^12^ Binding affinity of each peptide to the 67 HLA alleles were calculated by the NetMHCpan-4.0 algorithm^13^. The fraction of peptides bound by each HLA allele was determined for each dataset, using an established 500 nM affinity (IC50) cut-off for binding. The stability of complexes between HLA alleles and bound neopeptides was determined using the NetMHCstabpan-1.0 algorithm^14^, using the option of excluding prior information on exact binding affinity values (IC50). Half-life of peptide-HLA complex in hours was used as a proxy for peptide-HLA complex stability.

### Calculating peptidome diversity from mass spectrometry studies

The accuracy of the Pr estimates was tested using the mass spectrometry immunopeptidomes derived from mono-allelic cell lines^15,16^. For each HLA class-I allele with appropriate peptidome data, sequence diversity of the identified peptides was measured by the Shannon entropy index.

### Analysis of cancer patients treated with ICI therapy

Data on published cohorts of patients were collected who had melanoma treated with CTLA-4 inhibitors^17–19^, melanoma treated with PD-1 inhibitors^18^ and NSCLC treated with PD-1 inhibitors^20^. Information on HLA genotype, loss of HLA heterozygosity and cancer mutation burden in these patients were derived from Ref ^18^. Genotype-level HLA promiscuity (or genotype Pr) was determined by calculating the mean of 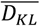 values across all HLA-A, B and C alleles of the patient and taking its reciprocal value. For patients with a single HLA-allele with no appropriate allelic Pr data, a standard imputation technique was involved by replacing the missing Pr with the median of that variable for all other cases. This protocol has the benefit of not changing the sample median for Pr, and it enlarges the dataset, but reassuringly, it has no impact on the association between survival and Pr (data not shown). Patients having at least two HLA alleles with unknown allelic Pr were excluded from the analysis.

Data on HLA-I evolutionary divergence (HED) was derived from ref ^21^. Survminer^22^ and Survival^23^ R libraries were used for statistical analysis of patient survival. The effect of genotype Pr on survival was tested with logrank test and by the ggsurvplot function of Survminer R library^22^, and the results were visualized by Kaplan-Meier curves. Survival curves of cohorts classified into more than two groups were also tested for ordered differences as implemented by the Survminer R library^22^. Cox regression models were generated using the coxph function of the Survival^23^ R library. The proportional hazard assumption for the Cox regression model fit was tested with the cox.zph function. The forest plot for multivariate cox models was generated with the forestmodel^24^ R library.

### Calculating differential agretopicity index (DAI)

Using literature information^5,7–11^, we compiled a dataset of experimentally verified neopeptides, all of which are known to have high binding affinity to certain HLA alleles (Supplementary data). To reach the best prediction accuracy with NetMHCpan-4.0 algorithm^13,25^, the analysis focused on nine amino acid long peptides. As earlier^6^, neopeptides with more than 50% sequence identity were excluded, resulting in 589 peptide neopeptides. Using an established method^26^, for each neopeptide, the peptide sequence was aligned against the non-mutated human proteome using BLAST+ 2.10.0^27^. The closest hit was retrieved, and the BLOSUM62 sequence similarities were calculated using an established method^28^.

The binding affinity of each neopeptide and the corresponding non-mutated human peptide to each of the 67 HLA-I alleles was calculated using NetMHCpan-4.0^13^. For each HLA-I peptide pair, DAI values were calculated as earlier^25^. For each HLA allele, the median DAI value was calculated across all neo-peptides with significant binding affinity to the given HLA variant (i.e. lower than 2% predicted binding rank percentile by NetMHCpan-4.0). The same method was used for calculating DAI values for 1030 immunogenic nine amino acid long viral epitopes acquired from ref^12^ (Supplementary Data).

DAI values were also determined for neopeptides in 139 melanoma samples derived from two immunotherapy cohorts^17,19^. Genomic data on missense mutations in protein coding genes were acquired from cBioPortal^29^. The corresponding mutated proteins were identified using the reference human proteome (downloaded from UniProt database^2^, as of 9^th^ January, 2020). Using these protein data, all possible 8 to 12 amino acid long mutated peptide segments were determined in each sample. Binding affinity and DAI values of each mutated peptide to each HLA class I allele of the patient was calculated as above.

### Transcriptome analysis of melanoma cancer samples

Normalized RNA-seq expression data of cancer samples derived from anti CTLA-4-treated melanoma patients^19^ with established HLA-I genotype were downloaded from cBioPortal^29^. Immune deconvolution analysis was carried out following recommendations by Sturm and colleagues^30^, and using the immunedeconv R library^30^. As recommended^30^, we used the EPIC algorithm^31^ for endothelial cells and cancer associated fibroblasts, MCP-counter^32^ for macrophages/monocytes and quanTIseq^33^ for regulatory CD4^+^ T cells. Precalculated TIDE dysfunction and exclusion score values were acquired from TIDE server^34^. Gene set enrichment analysis was carried out with the GSEA 4.0.3.^35^ software using 1000 permutations, weighted enrichment statistic and excluding gene sets with less than 5 genes. Gene sets of biological processes as reported by the Gene Ontology Consortium^36^ were included in the analysis. Gene sets enriched with P < 0.001 and FDR < 0.1 were treated as significant.

### Statistical analysis and visualization

Statistical analyses and visualization were carried out using R v.3.6.3^37^ under RStudio 1.2.5033 environment. Smooth curves on plots were fitted with cubic smoothing spline method^38^.

## Supplementary Material

**Figure S1.**
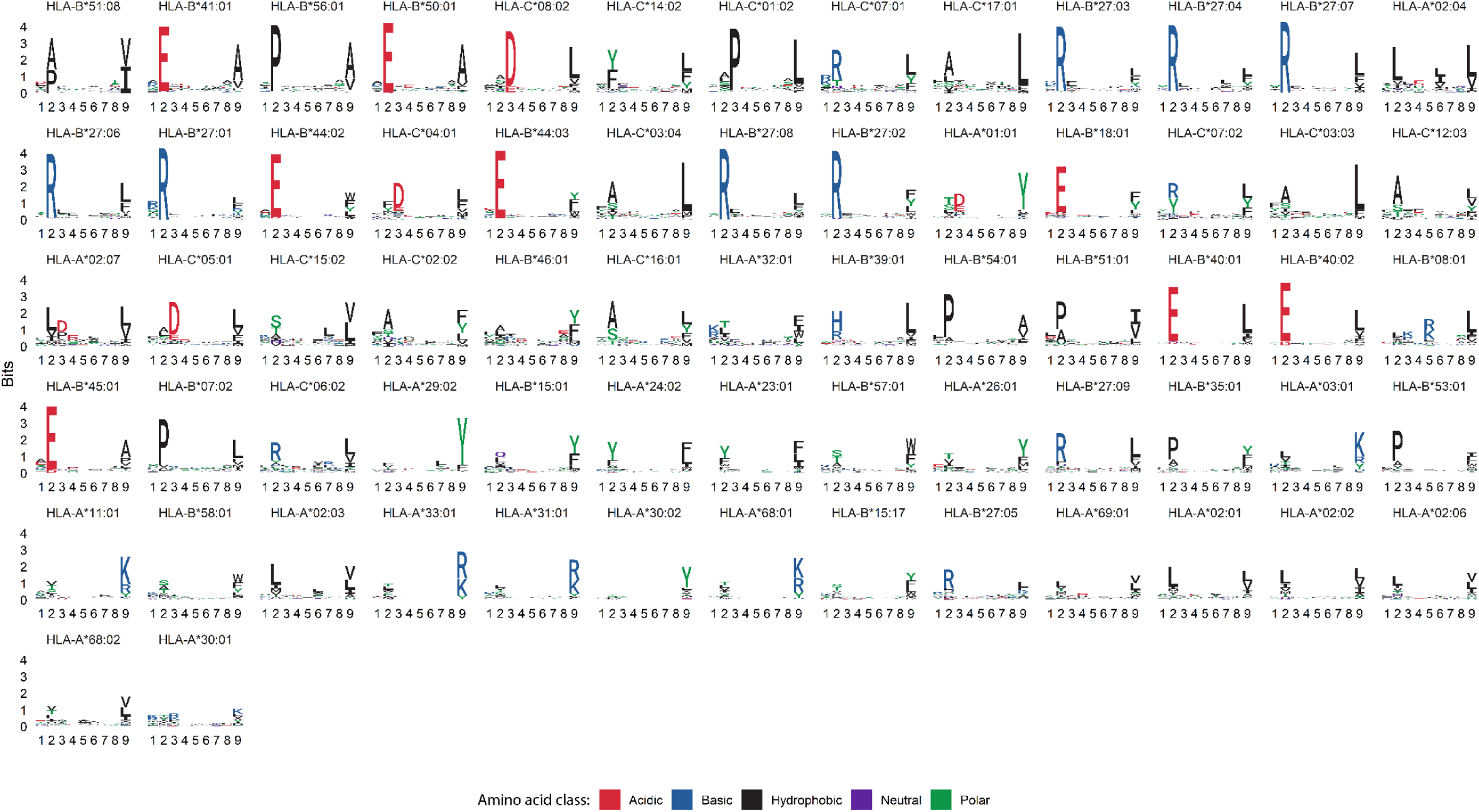
Sequence diversity of peptides presented by HLA class I alleles. Sequence diversity was calculated based on nine amino acid-long peptides bound by each of the 67 investigated HLA class I alleles and visualized as sequence logos^56^. Alleles are ordered according to their promiscuity level.

**Figure S2.**
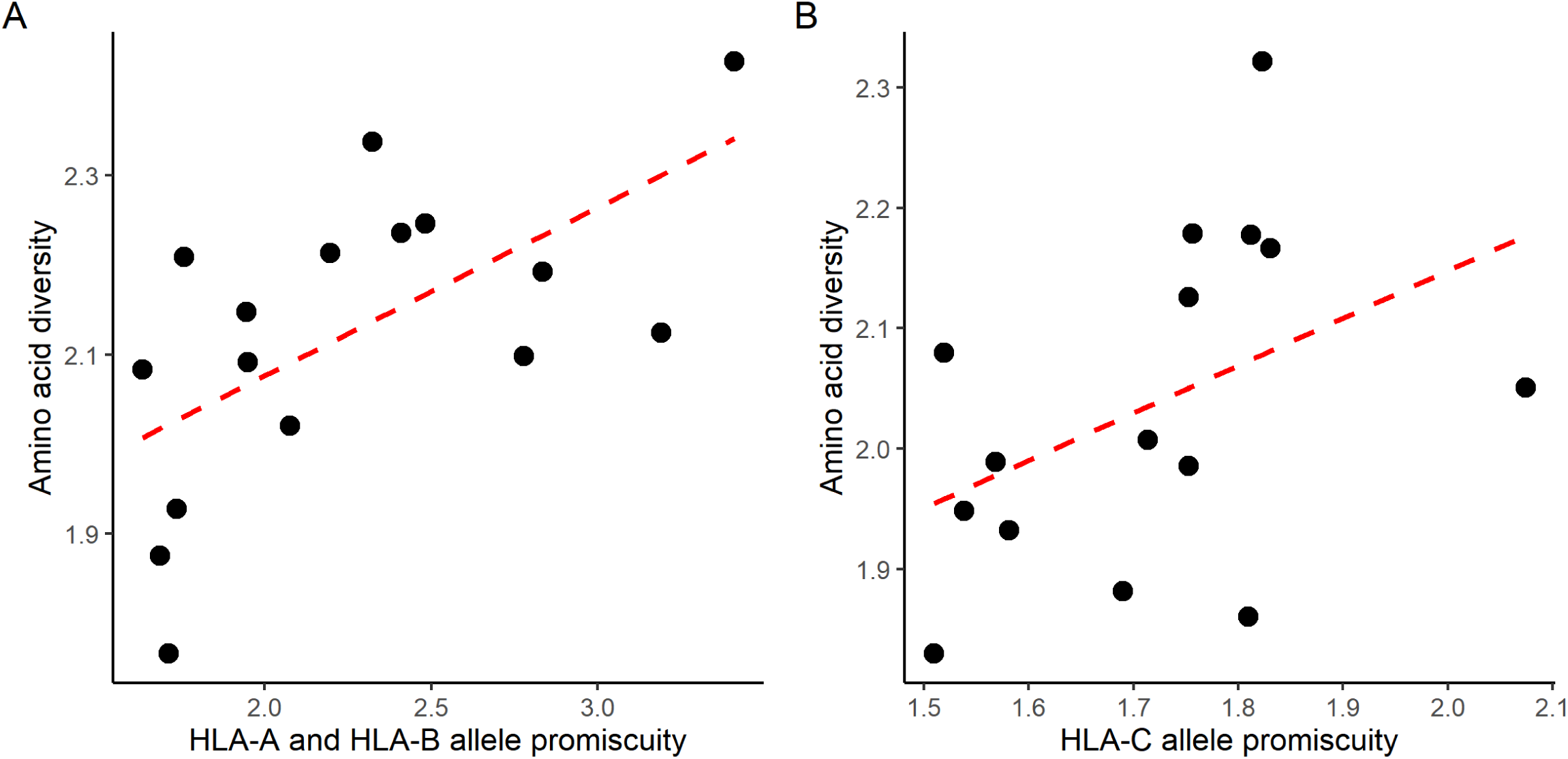
The relationship between HLA allele promiscuity and the diversity of presented peptides holds for two independent immunopeptidomics studies. We found a strong correlation between the promiscuity of alleles and sequence diversity of peptides eluted from the surface of monoallelic cell lines in studies by A) Abelin et al.^14^ and B) Di Marco et al.^15^ (Spearman’s rho: 0.67 and 0.56, P = 0.006 and 0.034 for panels A and B, respectively). Dashed red lines indicate linear regression line.

**Figure S3.**
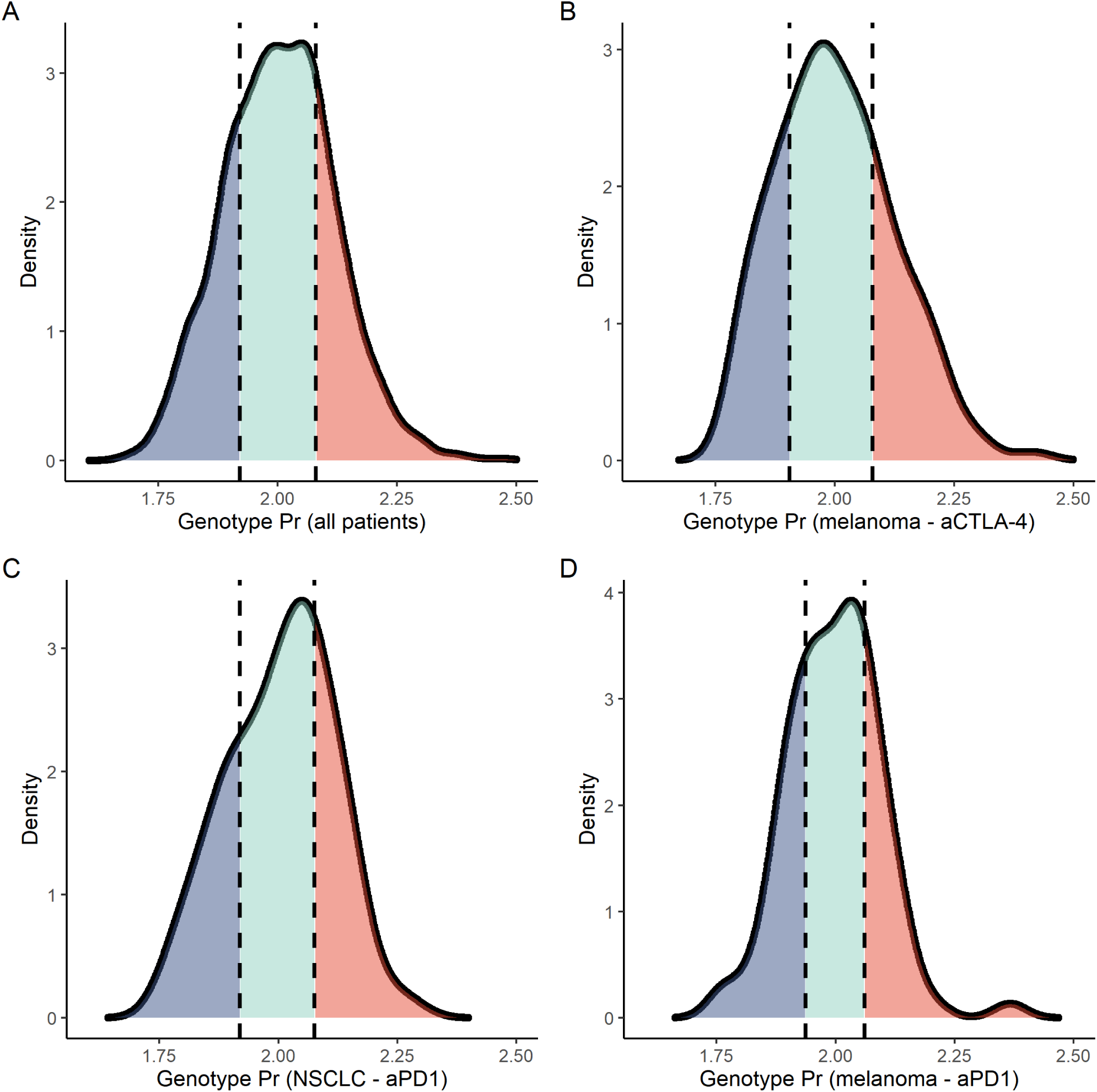
Distribution of genotype Pr in cancer immunotherapy cohorts. The density of genotype Pr is shown for A) all patients, B) melanoma patients treated with CTLA-4 inhibitors, C) non-small cell lung cancer (NSCLC) patients treated with PD1 or PDL-1 inhibitors and D) melanoma patients treated with PD1 or PDL-1 inhibitors. Dashed vertical lines represent the 25^th^ and 75^th^ percentile values.

**Figure S4.**
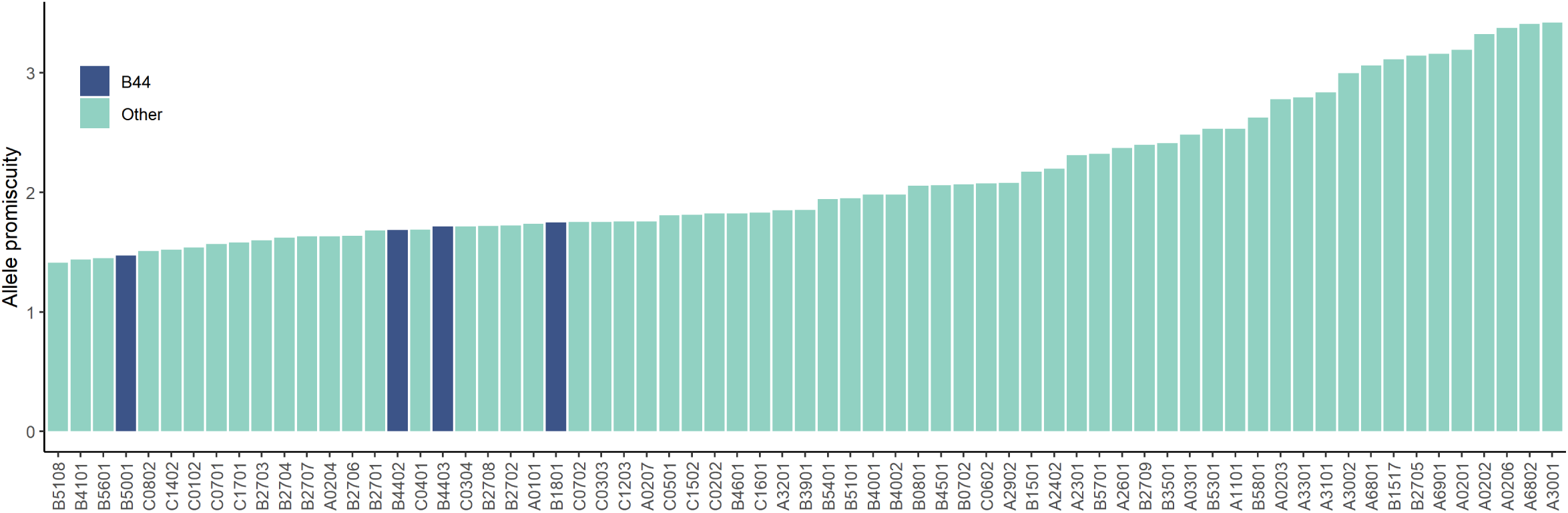
HLA alleles belonging to the B44 supergroup have low promiscuity. B44 alleles are highlighted in blue, and have significantly lower promiscuity than alleles not belonging to this group (Wilcoxon’s rank-sum test P: 0.049).

**Figure S5.**
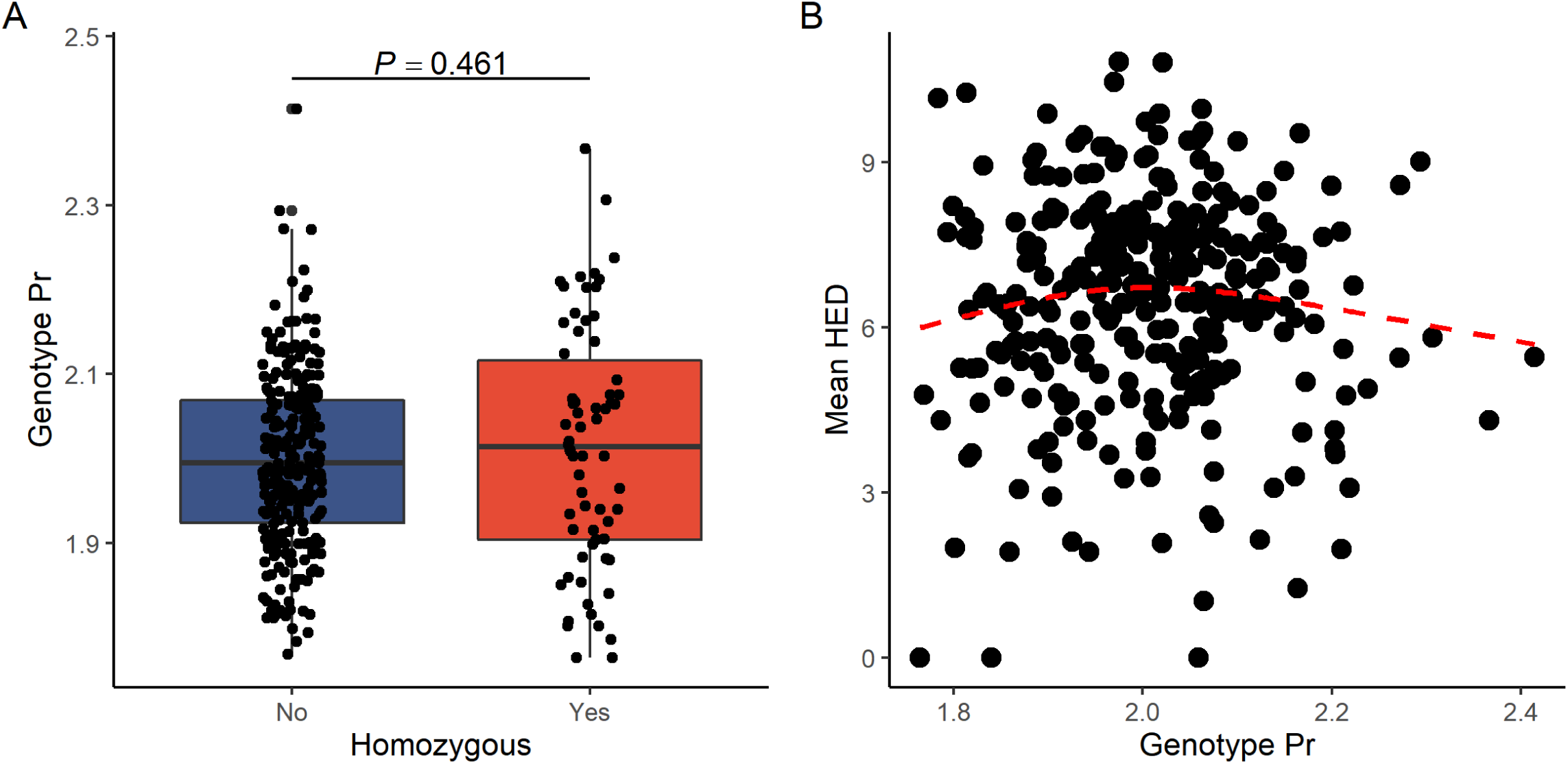
No relationship between genotype Pr, HLA homozygosity and HED. A) No significant difference in genotype Pr was found between patients fully HLA heterozygous patients and ones homozygous for at least one HLA-I locus. The P-value for Wilcoxon’s rank sum test is indicated. B) Similarly, there was no relationship between genotype Pr and mean HED of cancer immunotherapy patients (Spearman’s rho: -0.01, P = 0.79). Dashed red line indicates a smooth curve fitted using cubic smoothing spline method in R (see Methods). Patients from Fig. 3 A to C were included in both analyses. On boxplot, horizontal lines indicate median, boxes indicate the interquartile range (IQR), vertical lines indicate 1^st^ quartile - 1.5*IQR and 3^rd^ quartile + 1.5*IQR.

**Figure S6.**
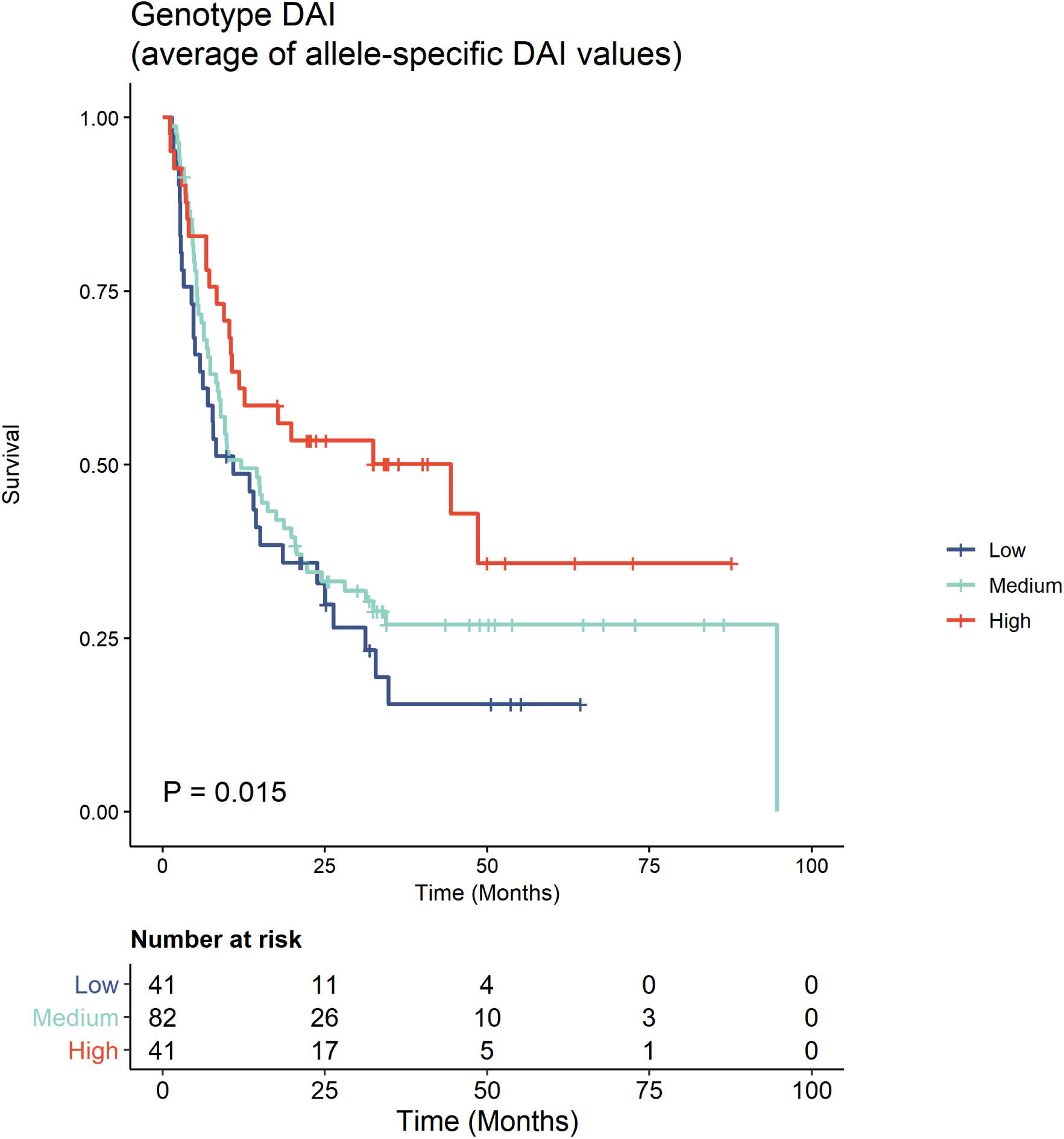
The effect of genotype DAI on survival when determined as the average of allele-specific DAI values. Patients were stratified into groups using the 25^th^ and 75^th^ percentile of all values as cutoffs. Log-rank test P-value is shown. Between-group differences were also tested for trend (see Methods). Higher genotype DAI value was associated with better survival.

**Figure S7.**
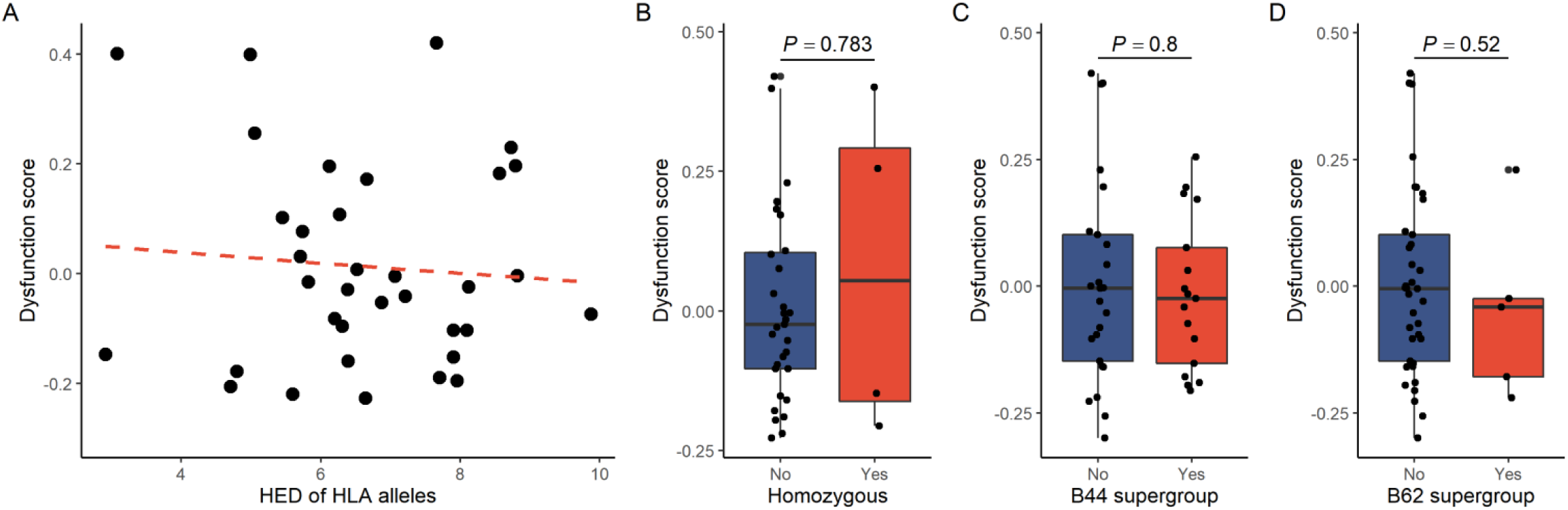
The effect of HLA-associated features on TIDE dysfunction score. A) The TIDE dysfunction score is shown as a function of HLA evolutionary divergence. There was no relationship between the two variables (Spearman’s rho: -0.01, P = 0.96). Similarly, neither HLA homozygosity (B), nor the carrier status for B44 (C) or B62 (D) supertype alleles affected the TIDE dysfunction score. P-values indicate Wilcoxon’s rank sum test results. On plot A, dashed red line indicates a linear regression line. On boxplots, horizontal lines indicate median, boxes indicate the interquartile range (IQR), vertical lines indicate 1^st^ quartile - 1.5*IQR and 3^rd^ quartile + 1.5*IQR.

**Figure S8.**
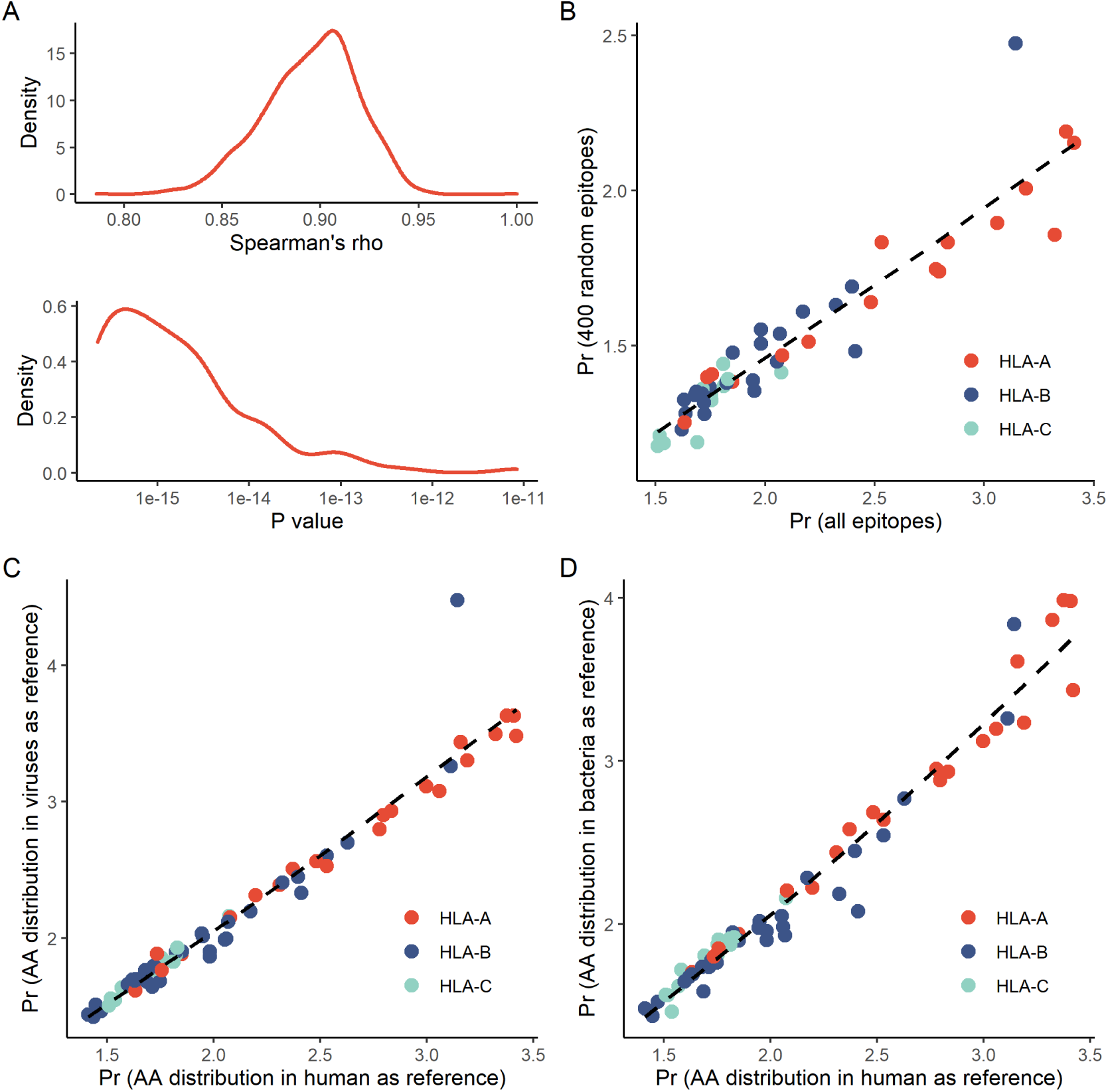
Promiscuity can be reliably inferred using a minimum threshold of 400 peptide sequences per allele. A-B) The number of identified peptide sequences for a given HLA allele is variable, which may have a significant effect on the calculated Kullback-Leibler (KL) divergence values. To test whether KL divergence can be reliably determined using a minimal peptide sequence number of 400, we first selected HLA alleles with large (more than 1000) number identified peptides (N = 51). After randomly selecting 400 sequences from the repertoire of each allele, we calculated KL divergence as explained in the Methods section. We iterated this process for 1000 times and compared the Pr values with those calculated using the full dataset. Panel A indicates the distribution of Spearman’s rho values and the P-values for correlations. B) An example plot indicating the relationship between promiscuity values calculated using either 400 random or all peptides bound by a given allele. C-D) The calculated promiscuity values are independent of the reference amino acid (AA) distribution used during calculation. The Pr values showed strong a correlation when calculated using C) human and viral or D) human and bacterial AA distributions as a reference (Spearman’s rho: 0.99, P < 2.2*10^-16^ for both comparisons). On plots B to D, dashed lines indicate a smooth curve fitted using cubic smoothing spline method in R (see Methods)

**Table S1.** The effect of HLA allele promiscuity on diversity and the fraction of bound peptides is independent of HLA loci. Results for multivariate models including allele promiscuity as continuous and HLA locus as categorical independent variables are shown. “P (var.)” values indicate the probability of observing a relationship between promiscuity and patient survival due to chance. Adjusted R-squared and F-test P-value of each model is indicated. Only the coefficient for allele promiscuity is shown.

| Dependent variable | Coeff. | P (var.) | Adjusted R-squared | F-test P-value |
| --- | --- | --- | --- | --- |
| Sequence diversity on cell surface | 0.203 | 0.004 | 0.275 | 0.008 |
| Fraction of neopeptides bound | 1.136 | 0.004 | 0.417 | 4.3 * 10<sup>-08</sup> |
| Fraction of human peptides bound | 1.205 | 9.6 * 10<sup>-05</sup> | 0.264 | 5.3 * 10<sup>-05</sup> |
| Fraction of viral peptides bound | 0.035 | 0.006 | 0.408 | 6.6 * 10<sup>-08</sup> |

**Table S2.** The effect of HLA allele promiscuity on survival is independent of the effect of a single allele. We analyzed the enrichment of alleles in different promiscuity groups of the merged cohort including all patients from Figure 3A to C (N = 327). Only those alleles were included in the analysis, which were carried by at least 10% of the individuals. We stratified patients into low, medium and high genotype Pr groups based on the 25^th^ and 75^th^ rank percentiles of all values. Next, we determined the odds ratio (OR) for finding a given allele in each promiscuity group and calculated if the enrichment is significant using Fisher’s exact tests. Significant FDR-corrected P-values are highlighted in red. Alleles enriched in low or high promiscuity groups are highlighted in blue or red background colours, respectively. Next, we excluded individuals carrying the given allele from the cohort, and determined the effect of promiscuity (as a continuous variable) on survival using Cox models. The effect and statistical significance of HLA promiscuity, and the Log-rank test P for the model is shown in the table. P-values lower than 0.05 are highlighted in red.

| Allele | OR in low | OR in medium | OR in high | Fisher's exact test P | Fisher's exact test FDR | Num. of patients when excluded | Coeff. for prom. | P (prom.) | Log-rank test P |
| --- | --- | --- | --- | --- | --- | --- | --- | --- | --- |
| A0201 | 0.221 | 0.994 | 3.774 | $1.6 * 10^{-14}$ | $3.6 * 10^{-13}$ | 175 | 2.405 | 0.025 | 0.030 |
| A0101 | 3.556 | 0.832 | 0.210 | $7.5 * 10^{-13}$ | $8.7 * 10^{-12}$ | 225 | 2.860 | 0.002 | 0.002 |
| B0801 | 3.541 | 0.675 | 0.164 | $2.1 * 10^{-11}$ | $1.6 * 10^{-10}$ | 264 | 2.599 | 0.001 | 0.001 |
| C0701 | 3.089 | 0.898 | 0.213 | $3.6 * 10^{-11}$ | $2.1 * 10^{-10}$ | 226 | 2.751 | $8.8 * 10^{-04}$ | 0.001 |
| B3501 | 0.173 | 0.996 | 2.032 | $2.5 * 10^{-04}$ | 0.001 | 285 | 3.126 | $6.7 * 10^{-05}$ | $8.9 * 10^{-05}$ |
| C0401 | 0.460 | 0.946 | 1.858 | 0.001 | 0.004 | 250 | 2.780 | $9 * 10^{-04}$ | 0.001 |
| A2402 | 0.912 | 1.479 | 0.305 | 0.001 | 0.004 | 270 | 2.411 | 0.001 | 0.002 |
| C0602 | 0.492 | 0.801 | 2.124 | 0.001 | 0.004 | 274 | 2.479 | 0.005 | 0.006 |
| C0202 | 0.116 | 1.281 | 1.419 | 0.004 | 0.010 | 295 | 2.015 | 0.008 | 0.009 |
| B4403 | 1.877 | 1.030 | 0.221 | 0.004 | 0.010 | 294 | 2.134 | 0.004 | 0.005 |
| B1501 | 0.386 | 0.910 | 1.918 | 0.008 | 0.017 | 288 | 2.047 | 0.007 | 0.008 |
| B1801 | 1.743 | 0.938 | 0.490 | 0.044 | 0.084 | 289 | 2.128 | 0.004 | 0.005 |
| C0304 | 0.595 | 0.854 | 1.778 | 0.054 | 0.096 | 288 | 2.098 | 0.006 | 0.007 |
| B4402 | 1.781 | 0.776 | 0.770 | 0.067 | 0.111 | 287 | 2.173 | 0.003 | 0.004 |
| C0702 | 0.566 | 1.181 | 1.137 | 0.084 | 0.129 | 244 | 2.256 | 0.003 | 0.004 |
| B5101 | 0.475 | 1.291 | 0.990 | 0.122 | 0.175 | 287 | 2.346 | 0.002 | 0.002 |
| C0501 | 1.596 | 0.827 | 0.822 | 0.152 | 0.206 | 280 | 2.205 | 0.003 | 0.004 |
| B0702 | 0.614 | 1.135 | 1.161 | 0.167 | 0.214 | 249 | 2.191 | 0.005 | 0.006 |
| A1101 | 0.595 | 1.266 | 0.904 | 0.271 | 0.328 | 288 | 2.112 | 0.005 | 0.005 |
| A0301 | 0.777 | 1.011 | 1.225 | 0.439 | 0.505 | 234 | 2.495 | 0.002 | 0.002 |
| C0303 | 0.610 | 1.137 | 1.129 | 0.506 | 0.555 | 295 | 2.249 | 0.003 | 0.003 |
| C1203 | 0.903 | 1.171 | 0.770 | 0.617 | 0.645 | 287 | 2.210 | 0.004 | 0.004 |
| A2601 | 0.951 | 1.062 | 0.925 | 0.972 | 0.972 | 293 | 2.213 | 0.003 | 0.003 |

**Table S3.** The effect of genotype Pr on patient survival when calculated using individual loci. We examined four univariate Cox models containing the mean promiscuity of individual loci (Models 2 to 4) and all loci (Model 1) as predictors. The effect of variables on patient survival is shown. “P (var.)” values indicate the probability of observing a relationship between the predictor variable and patient survival due to chance. Log-rank test P-values and the log-likelihood values for individual models are also indicated. Significant ANOVA P-values indicate that the first model fits better than the examined one. NA: not applicable. P-values lower than 0.05 are highlighted in red.

| Model No | Locus | Coeff. | P (var.) | Log-rank test P | Log-likelihood | ANOVA P (vs. Model 1) |
| --- | --- | --- | --- | --- | --- | --- |
| 1 | A + B + C | 2.362 | 9.1 * 10<sup>-04</sup> | 0.001 | -706.750 | NA |
| 2 | A | 0.317 | 0.152 | 0.153 | -711.077 | < 2 * 10<sup>-16</sup> |
| 3 | B | 1.084 | 0.004 | 0.005 | -708.233 | < 2 * 10<sup>-16</sup> |
| 4 | C | 0.973 | 0.219 | 0.225 | -711.362 | < 2 * 10<sup>-16</sup> |

